# Seeking the Membrane-Bound Structure of the Caveolin 8S Complex

**DOI:** 10.1101/2025.03.09.642159

**Authors:** Sayyid Yobhel Vasquez Rodriguez, Themis Lazaridis

## Abstract

The protein caveolin-1 (CAV1) is essential in the generation of caveolae, cup-like invaginations in the plasma membrane, but the mechanism of its action remains unclear. A recent cryo-EM structure showed an 11-mer of CAV1 (the 8S complex) forming a disk with a flat membrane-facing surface, raising the question of how a flat complex is able to generate membrane curvature. We previously conducted implicit-solvent molecular dynamics simulations, which showed the 8S complex adopting a conical shape, with its outer ridge deep inside the implicit membrane. These results suggested a scaffolding-type mechanism for curvature generation by the 8S complex. In this work we aimed to validate this proposal via all-atom simulations. To date, all simulations (other than in vacuum) show the complex taking a conical shape. The arrangement of lipids around the complex depends on the starting configuration. Starting on top of the bilayer leads to lipid extraction and water molecules trapped between the 8S complex and the bilayer, creating a protrusion on the distal leaflet. Starting deep inside the bilayer, displacing the proximal leaflet, leads to a more plausible configuration with the distal leaflet lipids adsorbed onto the 8S concave surface. Further work is needed to characterize the determinants of 8S shape and its membrane curvature generating capabilities, as well as the role of lipid composition.

## INTRODUCTION

Caveolae, small cup-like invaginations of the plasma membrane prevalent in mammalian cells ^1^, perform multiple cellular functions. They protect the plasma membrane from mechanical damage by disassembling and flattening, which increases the plasma membrane’s surface area and allows the cell to regulate membrane tension. They are suggested to indirectly play a role in the cell’s endocytic processes and perform a specialized form of endocytosis in blood cells. Lastly, caveolae regulate the plasma membrane’s lipid composition and its signaling pathways. The importance of these functions is underscored by disease phenotypes such as muscular dystrophies, lipodystrophy, and various cancers linked to the dysfunction of caveolae ^1–3^. Caveolae are formed primarily by the coordinated actions of three caveolin and four cavin proteins ^1^. The present study focuses on the 8S complex, an oligomeric form of caveolin-1 (CAV1).

CAV1 is primarily expressed in adipocytes, fibroblasts, endothelial cells, smooth muscle cells, and type 1 pneumocytes ^4,5^. It is composed of 178 amino acid residues divided into the following domains: N-terminus (1-81), scaffolding domain (SD, 82– 101), oligomerization domain (61-101), intramembrane domain (IM, 102–134), and C-terminus (135–178) ^6^. Caveolin expression by itself can deform the plasma membrane since it was shown to produce caveola-like vesicles in recombinant prokaryotic models ^7^. Furthermore, CAV1 was shown to produce wide pan-like invaginations on genetically engineered mouse embryonic cells ^8^.

CAV1 is classified as an integral membrane protein since it is co-translationally inserted into the endoplasmic reticulum before being incorporated into the plasma membrane, requires detergent for removal from the plasma membrane, and moves through multiple interior compartments for recycling purposes; all characteristics of integral membrane proteins ^9^. CAV1’s characteristics as an integral membrane and its topology, where both the N and C termini domains are exposed to the cytoplasm, led to the proposal that the IMD acts as a helical hairpin that wedges and deforms the plasma membrane ^10^. However, a new CAV1 8S complex cryo-EM structure exhibited characteristics that differed from the helical hairpin model and were more reminiscent of a peripheral protein. The CAV1 8S structure showed the IMD contributing to the formation of a flat membrane-facing surface and was proposed to stabilize the flat membrane faces of polyhedral structures instead of deforming the plasma membrane ^11^.

These findings called into question the manner by which is CAV1 able to bend the membrane in the absence of cavins. To answer this question, we previously conducted molecular dynamics simulations of the CAV1 8S complex (PDB: 7SC0) in an implicit solvent membrane ^12^. In these simulations, the CAV1 8S complex took a conical shape with a concave membrane-binding surface and its outer ridge buried deep within the membrane. Additionally, we found especially strong membrane binding in vesicles of 5-10 nm radii with protonation of the residue E140. The discrepancy from the experimental structure would normally lead to discounting the simulations. However, the simulated concave model makes more mechanistic sense and offers a clear mechanism for membrane curvature by caveolin proteins. Furthermore, we speculated that some experimental factor biases the experimental structures towards a flatter complex, such as the presence of large amounts of detergent used to segregate the complex ^12^.

One significant limitation of this previous work was that the lipids were implicit and the membranes idealized and non-deformable. Thus, one could not tell how the lipid molecules were arranged around the complex. The goal of the current project was to address this limitation by simulating the CAV1 8S complex in an all-atom lipid bilayer. To start, we chose a standard POPC bilayer and initially placed the complex at different depths within this bilayer. The results are compared to recently reported all-atom simulations of the CAV1 8S in POPC/cholesterol membranes ^13^, coarse-grained simulations ^14^, and a theoretical model for curvature generation by the flat cryoEM complex ^15^.

## METHODS

The all-atom systems were created in the CHARMM-GUI server ^16^ based on the cryo-EM structure (PDB 7SC0) ^11^. One system consisted of the complex in a water box (191X191X96 Å, about 363,000 atoms) and the rest were protein-membrane systems with 818 POPC molecules and unit cells of about 165X165X142 Å or 178X178X115 Å with about 400,000 atoms. The water thickness was set to 32 Å. The system was neutralized with potassium ions in addition to 0.15 M KCl. All systems used the TIP3P model for water ^17^ the charmm36m force field for proteins ^18^ and the charmm36 force field for lipids ^19^. The systems were equilibrated locally using NAMD following the standard CHARMM-GUI protocol and then subjected to simulations on the Anton 2 supercomputer.

For the first set of simulations, five systems were created: four protein-membrane systems and one water box-protein system. In water the residue E140 was unprotonated. Because the protonation of E140 was found to enhance membrane binding in our previous implicit membrane simulations ^12^, half of the protein-membrane systems had the E140 residues protonated (E140H). The systems differed in the depth of placement of the 8S complex in the membrane. The starting structure was set with the principal axes of the complex along the x and y axis (the disk parallel to the xy plane). Then the complex was translated either 32 Å along the Z-axis, to place the 8S complex on top of the membrane, or 10 Å along the Z-axis, to embed the 8S in the membrane. In the latter placement the complex displaces completely the proximal leaflet. The systems were subjected to 1 μs simulations on the Anton 2 supercomputer. The 32 Å E140^-^ and E140H simulations were extended to 3 μs. The second set of simulations consisted of three all-atom protein-membrane systems using only unprotonated E140 residues with the 8S complex translated 16 Å, 25 Å, or 30 Å along the Z-axis. These systems were subjected to 0.25 μs simulations on the Anton 2 supercomputer. The 16-Å and 25-Å systems displace the proximal leaflet, while the 30-Å system does not. Table 1 shows the characteristics of the eight simulations performed.

**Table 1.** Number of lipids in each bilayer leaflet, number of water molecules and ions, and duration of the eight simulations. The membrane systems differ in the positioning of the complex with respect to the membrane center (designated by the amount of translation along the z axis).

| | Proximal | Distal | TIP3P | K<sup>+</sup>/Cl<sup>-</sup> | Duration (μs) |
| --- | --- | --- | --- | --- | --- |
| Water | - | - | 117691 | 366/333 | 1 |
| 32 Å (E140<sup>-</sup>) | 355 | 463 | 88951 | 278/245 | 3 |
| 32 Å (E140H) | 356 | 462 | 88887 | 267/245 | 3 |
| 10 Å (E140<sup>-</sup>) | 321 | 497 | 79812 | 252/219 | 1 |
| 10 Å (E140H) | 321 | 497 | 79768 | 241/219 | 1 |
| 16 Å (E140<sup>-</sup>) | 311 | 507 | 85126 | 271/238 | 0.25 |
| 25 Å (E140<sup>-</sup>) | 311 | 507 | 93436 | 296/263 | 0.25 |
| 30 Å (E140<sup>-</sup>) | 322 | 496 | 94498 | 299/266 | 0.25 |

Secondary structure was characterized using the DSSP method ^20^. As a quantitative method of determining the shape change in the 8S complex, we calculated the spatial extent (Å) of the complex (max – min x, y, or z coordinate) after it is placed with its principal axes on the xy plane.

## RESULTS

### The 8S complex becomes conical

As in our previous work ^12^, in all runs on Anton 2 the 8S complex changes shape from flat to conical. This includes the simulation in water (Fig. 1) and all membrane-bound systems (Figs. 2C-8C). The average spatial extent (Å) of the complex in all simulations is reported in Table 2. The extent of the complex along the Z axis (height) indicates how conical the 8S complex has become in comparison to the original flat shape. The water box system showed the largest height, followed by the systems translated furthest from the membrane. In the previous implicit simulations the extents were the same in water and somewhat larger on the membranes, especially the small vesicles (Table 3).

**Table 2.** Average spatial extent change (Å) of the 8S complex in the X, Y, and Z dimensions for each all-atom simulation. The standard deviations were between 1-4 Å.

| | $\Delta X$ | $\Delta Y$ | $\Delta Z$ |
| --- | --- | --- | --- |
| Cryo-EM structure | 137 | 137 | 40 |
| Water | 138 | 134 | 62 |
| 32 $\text{\AA}$ (E140$^-$) | 137 | 134 | 57 |
| 32 $\text{\AA}$ (E140H) | 136 | 132 | 56 |
| 10 $\text{\AA}$ (E140$^-$) | 134 | 132 | 54 |
| 10 $\text{\AA}$ (E140H) | 134 | 130 | 54 |
| 16 $\text{\AA}$ (E140$^-$) | 133 | 133 | 53 |
| 25 $\text{\AA}$ (E140$^-$) | 134 | 134 | 54 |
| 30 $\text{\AA}$ (E140$^-$) | 136 | 132 | 52 |

**Table 3.** Average spatial extent (Å) of the 8S complex in the Z direction for previously reported implicit solvent simulations ^12^. The standard deviations were between 1-4 Å.

| System | E140$^-$ | E140H |
| --- | --- | --- |
| Water | 62 | 60 |
| Flat membrane | 63 | 56 |
| 50-nm vesicle | 64 | 59 |
| 25-nm vesicle | 64 | 59 |
| 10-nm vesicle | 68 | 52 |
| 5-nm vesicle | 74 | 64 |

**Fig. 1.**
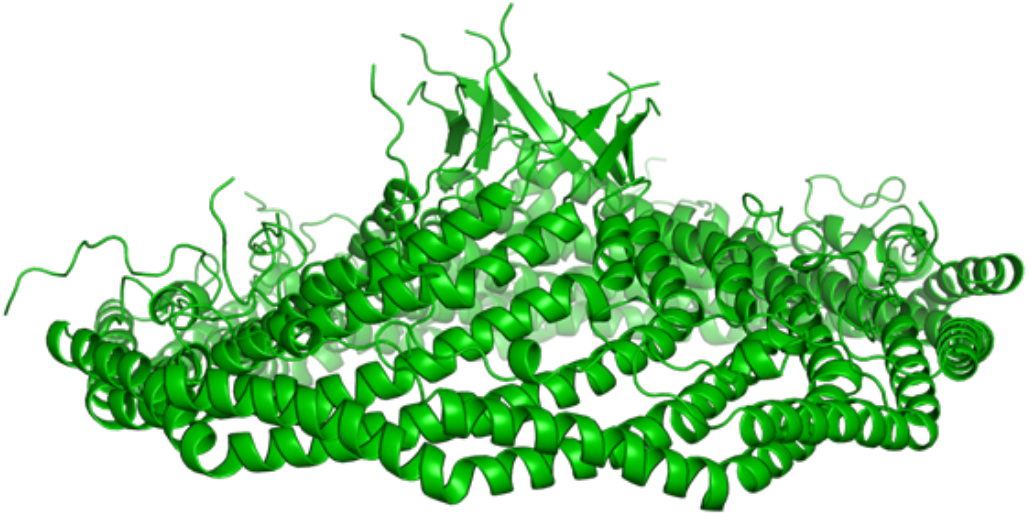
Final structure of the 1 μs simulation of the CAV1 8S complex in a water box. Protein-only lateral view.

**Fig. 2.**
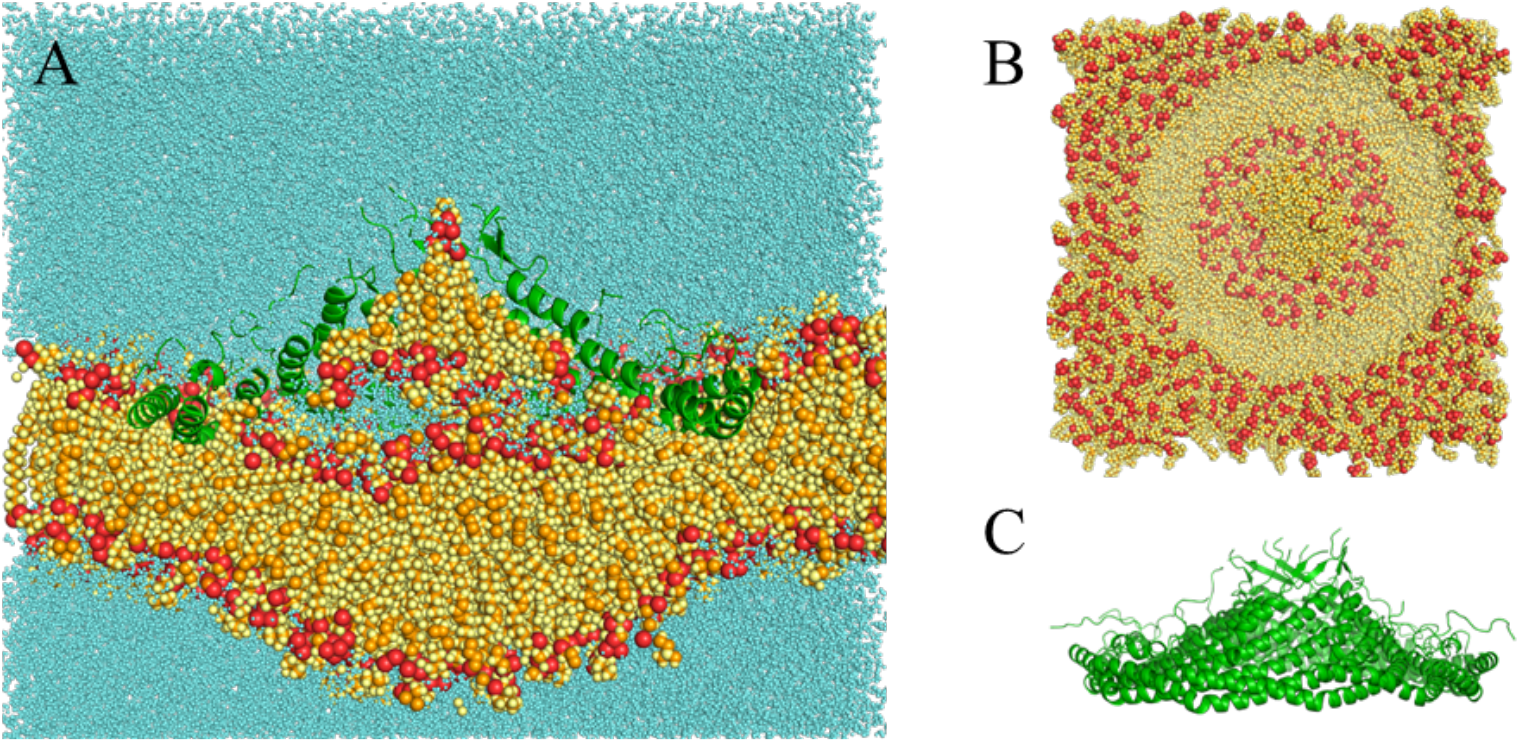
Final structure of the 3-μs simulation with E140^-^ at 32Å. (A) Complete system bisected lateral view. (B) Membrane-only top view. (C) Protein-only lateral view. The red particles are P and O, the orange particles are C, and all other atoms are yellow. The protein is represented as green ribbon.

Other than the shape change, the structure of the complex is well maintained, except for the beta barrel, which essentially dissolves in the water simulation and destabilizes to varying extents in the membrane simulations (Table 4). This is not surprising, as the interior of the barrel is hydrophobic. Helicity increases slightly in all simulations. The RMSD is largest for the N-terminal region, which takes different conformation in each protomer.

**Table 4.** Secondary structure and all-atom RMS deviations. Starting secondary structure is about 60-5-4-1 (% helix-beta-3_10_-π). The first RMS value is for the entire complex, the second for the beta barrel (residues 168-177), and the remaining for sections of the A protomer.

|  | 2ry str | RMS | barrel | 49-60 | 61-101 | 102-134 | 135-167 |
| --- | --- | --- | --- | --- | --- | --- | --- |
| Water | 63-2-1-1 | 7.7 | 7.3 | 6.2 | 2.6 | 1.7 | 2.8 |
| 32 Å (E140<sup>-</sup>) | 64-2-0-0 | 9.4 | 6.6 | 5.8 | 2.7 | 1.8 | 3.0 |
| 32 Å (E140H) | 63-3-2-1 | 8.3 | 10.8 | 5.5 | 2.3 | 1.8 | 3.3 |
| 10 Å (E140<sup>-</sup>) | 62-3-1-1 | 7.6 | 5.7 | 6.9 | 3.0 | 1.6 | 2.5 |
| 10 Å (E140H) | 63-2-1-1 | 7.1 | 5.5 | 5.9 | 2.1 | 1.8 | 2.3 |
| 16 Å (E140<sup>-</sup>) | 62-3-2-1 | 8.0 | 5.6 | 2.6 | 2.1 | 1.4 | 3.1 |
| 25 Å (E140<sup>-</sup>) | 62-2-2-1 | 9.3 | 5.9 | 4.7 | 1.9 | 1.4 | 2.6 |
| 30 Å (E140<sup>-</sup>) | 61-3-3-1 | 8.0 | 3.7 | 4.7 | 2.5 | 2.2 | 2.8 |

### Membrane-bound structure depends on starting configuration

For both 32-Å systems, the 8S complex pulled lipids from the proximal monolayer inward. In the deprotonated E140 system the extracted lipid cluster consists of about 20 lipids. Their orientation is with their headgroup down, toward the membrane, except for one lipid which pokes its headgroup into the barrel. In the E140H system only three lipids were extracted and they are oriented headgroup down or parallel to the membrane. At the same time, additional water entered the space between the complex and the membrane, forming a structure akin to a reverse micelle and resulting in a bulge on the other side (Fig. 2A & Fig. 3A). The water likely enters to solvate the zwitterionic lipid headgroups underneath the complex. It is not clear whether the difference in the number of lipids extracted is caused by the protonation of E140 or is a statistical fluctuation. E140H are seen to donate an HB to lipid ester or phosphate groups, whereas E140-interact primarily with positively charged choline. How this difference would favor lipid extraction is not immediately evident. Replicate runs in the future could resolve this question.

**Fig. 3.**
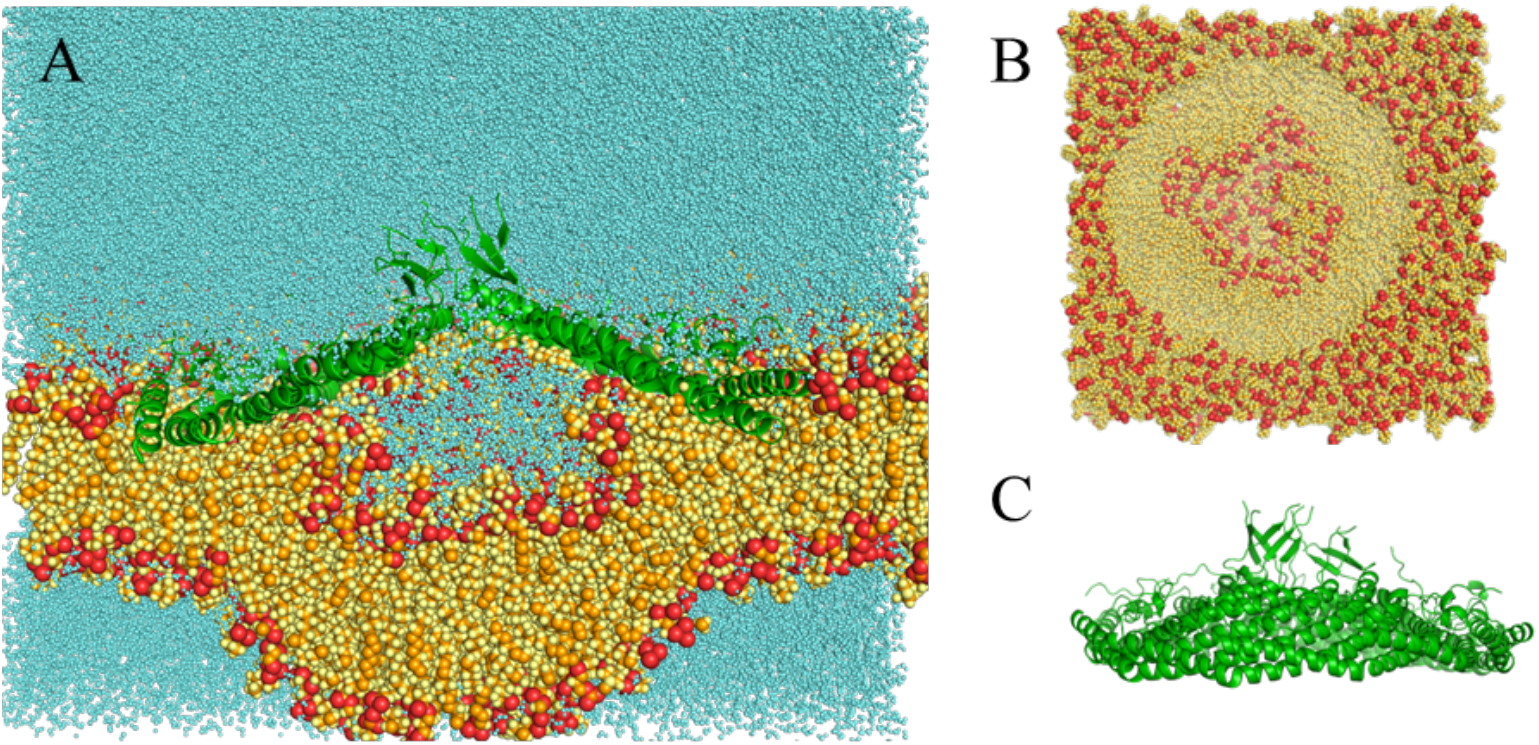
Final structure of the 3-μs simulation with E140H at 32 Å. (A) Complete system bisected lateral view. (B) Membrane-only top view. (C) Protein-only lateral view. Colors as in Fig. 2.

A very different picture arose when the complex was inserted deep into the membrane. For both 10-Å systems, the 8S complex pulled the distal monolayer toward itself (Fig. 4A & Fig. 5A). The coverage of the complex’s underside by lipid was not uniform, but exhibited distinct holes (Fig. 4B,5B). In the E140 system these holes appear near residue Tyr 151, which points toward the membrane. In the E140H system the holes are smaller and appear near A162. It is possible that these holes appear due to the limited availability of lipid in the distal leaflet and that they would repair given more time and additional amounts of lipid. The lack of cholesterol could also play a role (see Discussion).

**Fig. 4.**
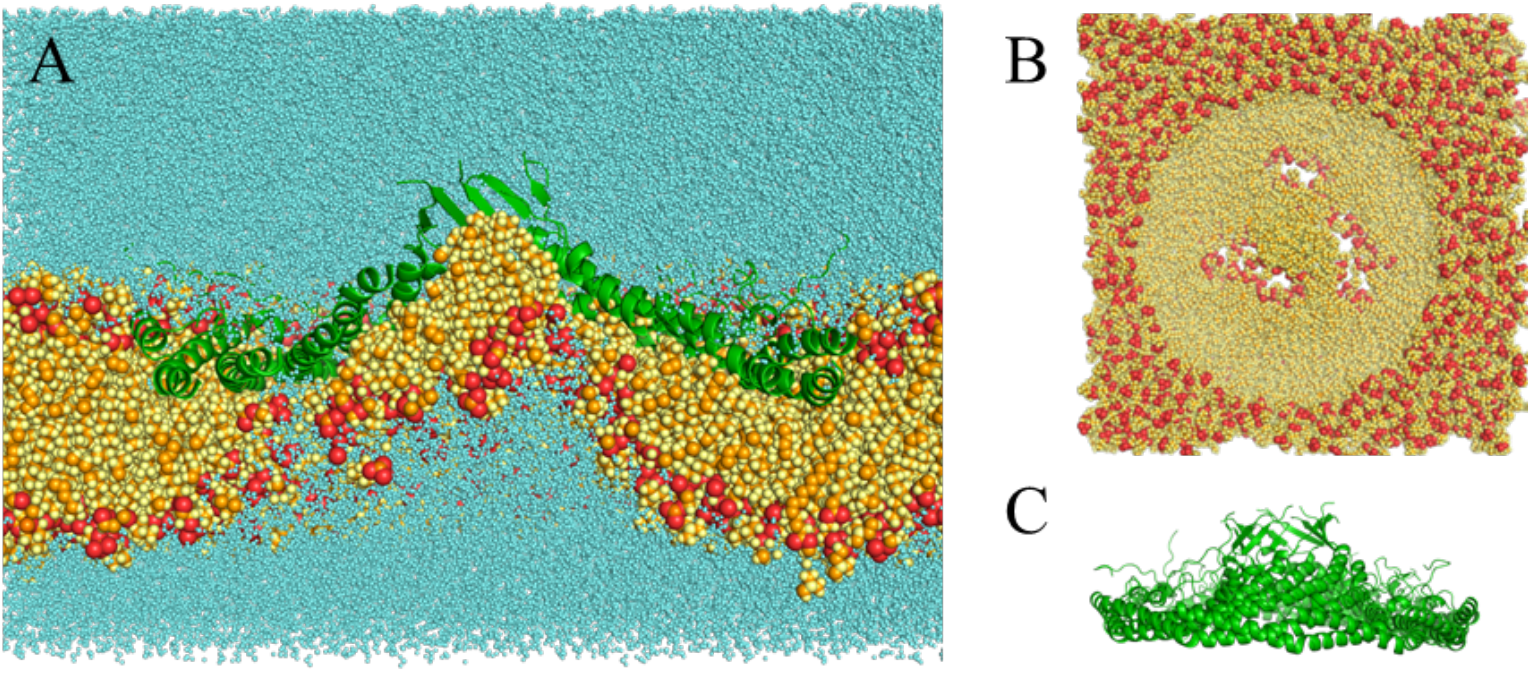
Final structure of the 1-μs simulation with E140-at 10Å. (A) Complete system bisected lateral view. (B) Membrane-only top view. (C) Protein-only lateral view. Colors as in Fig. 2.

**Fig. 5.**
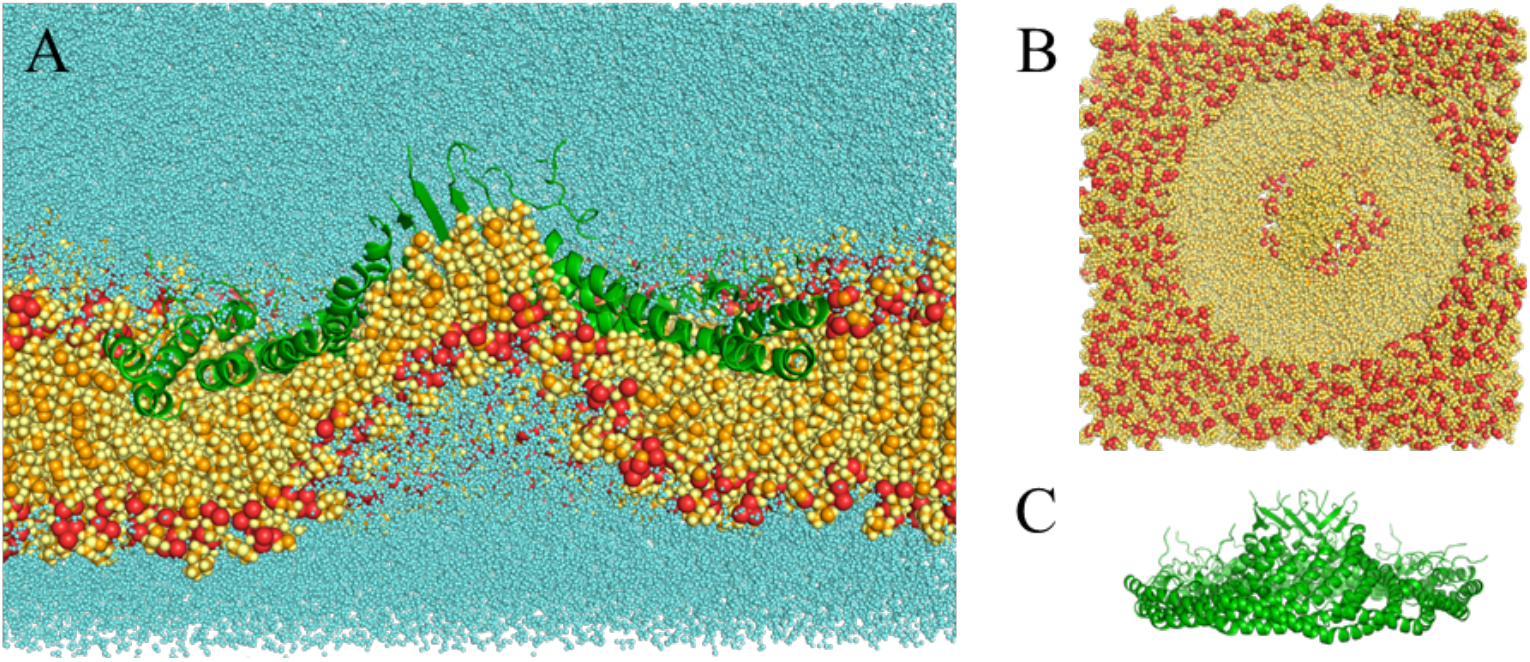
Final structure of the 1-μs simulation with E140H at 10Å. (A) Complete system bisected lateral view. (B) Membrane-only top view. (C) Protein-only lateral view. Colors as in Fig. 2.

For the 16-Å (Fig. 6) and 25-Å (Fig. 7) systems, the 8S complex pulled the distal monolayer inward, similar to the 10-Å systems from the first set. In the 30-Å system (Fig. 8), water was trapped between the complex and the membrane, pushing the membrane outwards, similar to the 32-Å systems from the first set. Here the beta barrel is filled with three lipids oriented with their headgroups down. As a result, this barrel exhibits the smallest RMS deviation from the starting structure (Table 4).

**Fig. 6.**
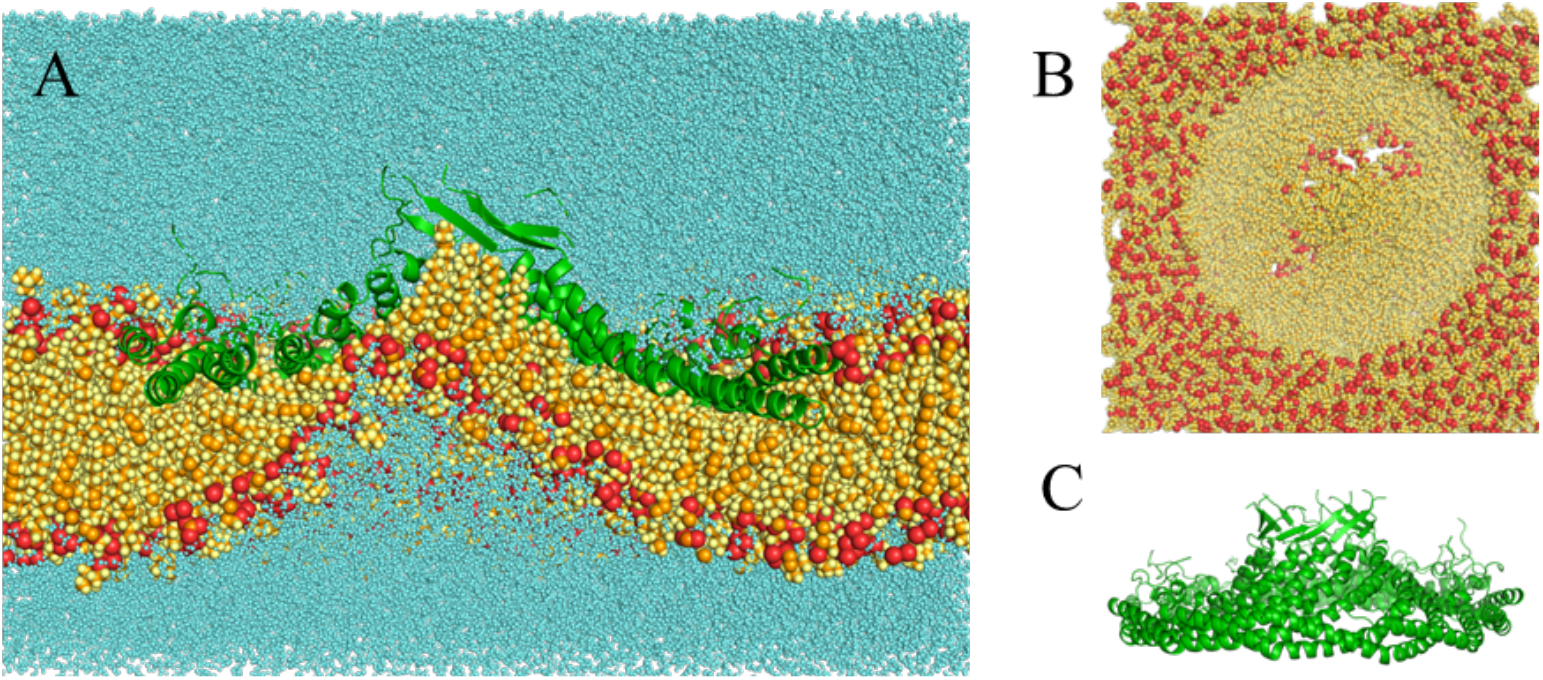
Final structure of the 0.25-μs simulation with E140^-^ at 16 Å. (A) Complete system bisected lateral view. (B) Membrane-only top view. (C) Protein-only lateral view. Colors as in Fig. 2.

**Fig. 7.**
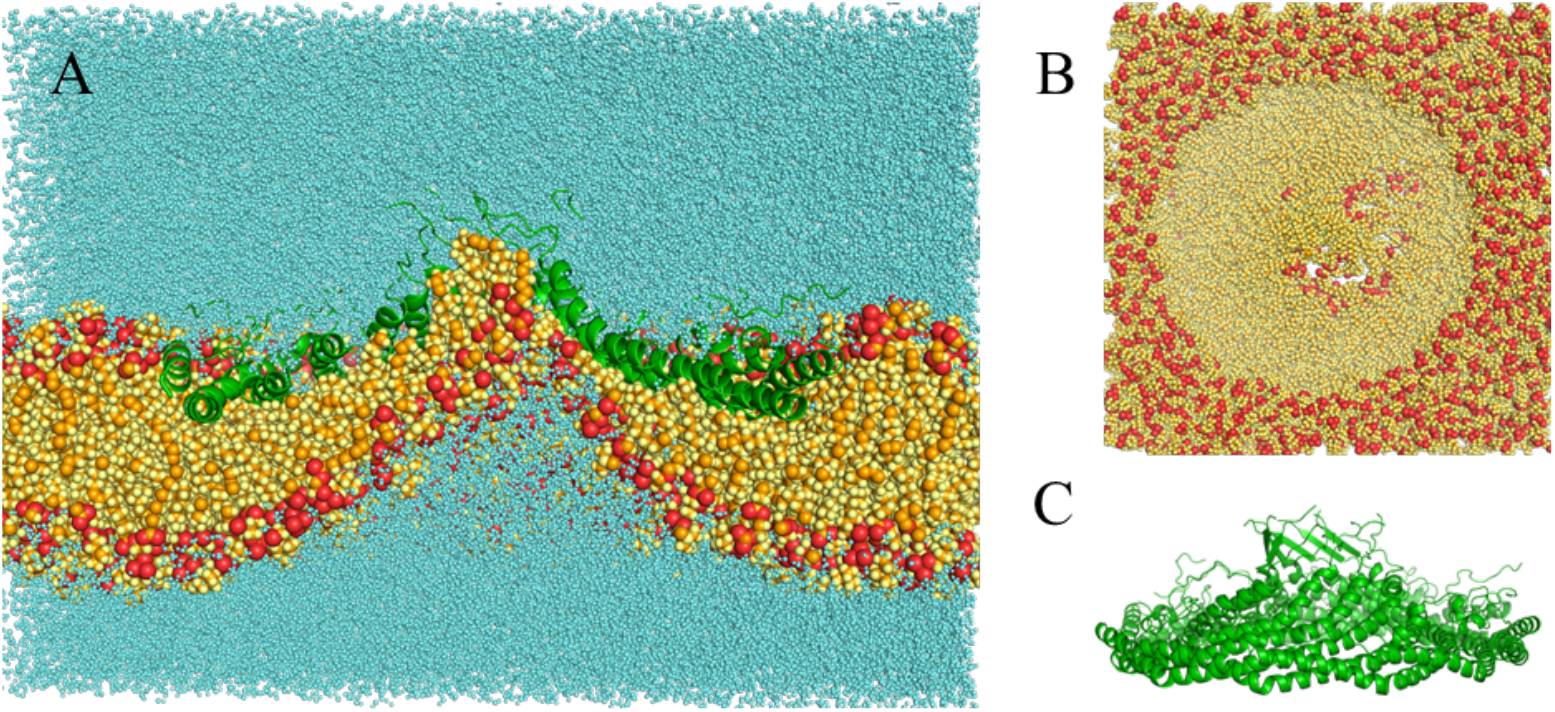
Final structure of the 0.25-μs simulation with E140^-^ at 25 Å. (A) Complete system bisected lateral view. (B) Membrane-only top view. (C) Protein-only lateral view. Colors as in Fig. 2.

**Fig. 8.**
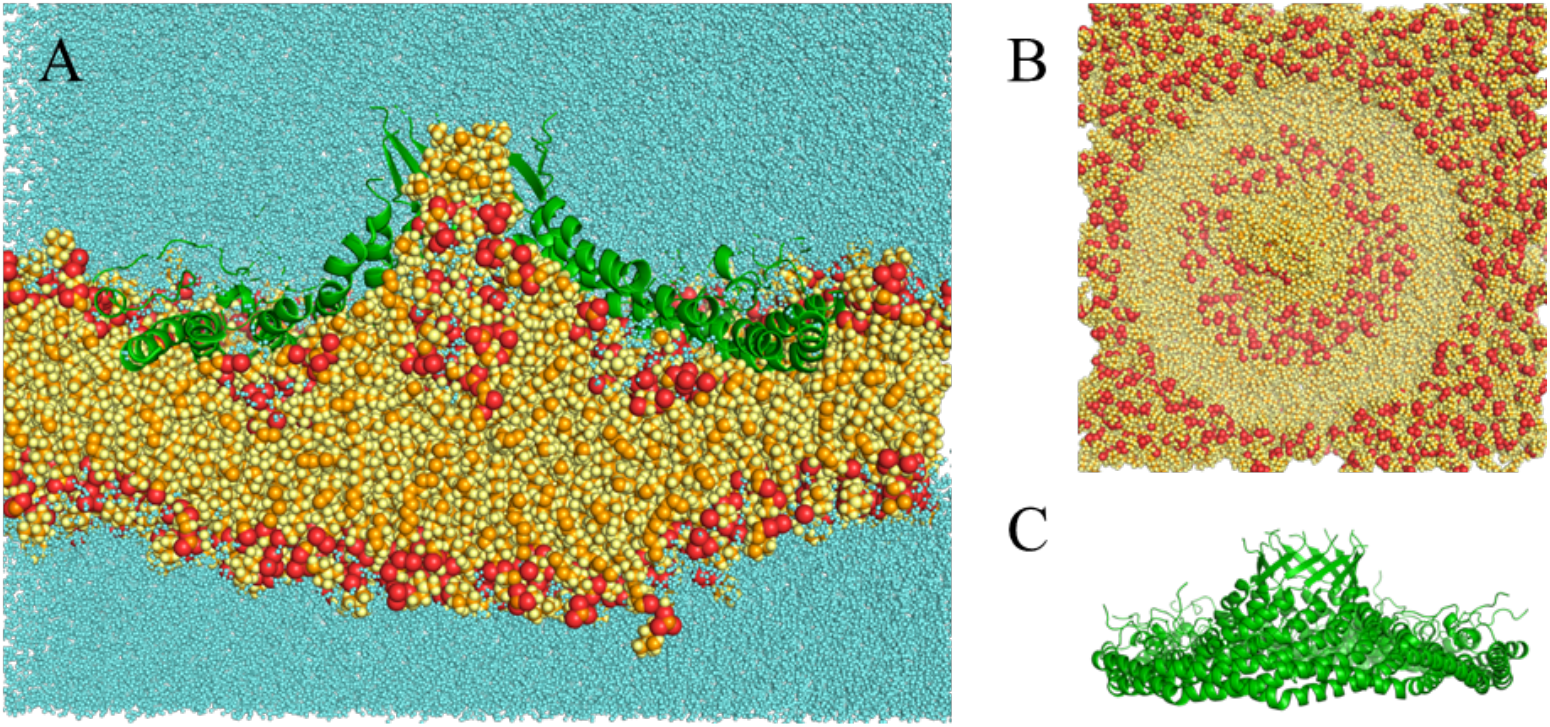
Final structure of the 0.25-μs simulation with E140^-^ at 30 Å. (A) Complete system bisected lateral view. (B) Membrane-only top view. (C) Protein-only lateral view. Colors as in Fig. 2.

### Origin of the shape change and possible role of the missing N-terminal region

The only simulation that was able to maintain the flat shape of 8S was one in vacuum. A 9-ns simulation of 8S in the charmm36 force field without explicit or implicit solvent gave a Z extent of 43.7 Å, only slightly higher than in the cryoEM structure. This small increase is not caused by a rise of the barrel but by a tilt of the barrel toward one side leading to a small bulge from the bottom of the disk. The conical conformation simulated in vacuum rapidly reverted to flat. This shows that the flat state is not kinetically trapped but thermodynamically more stable. Indeed, the energy of the flat conformation is lower than that of the conical one, favored by better van der Waals and bonded interactions. It has not been possible to pinpoint a small set of interresidue interactions responsible for the lower energy of the flat conformation in vacuum.

Another difference between the simulated and experimental system is in the 48 N-terminal residues (Nt region), which are present but invisible in the experiment and absent in the simulations. Alphafold2 predicts, with low confidence, a helix and a beta hairpin in this region (entry Q03135). Two independent 1-ns simulations in implicit solvent after adding the AF model for the Nt region to the cryoEM structure resulted in a structure that was less conical than those without the Nt region (height 49-52 Å vs. 40.1 Å in the cryoEM structure and ∼60 Å in the truncated system). The Nt regions in the final structure adopt different configurations and some of them interact with the side or the top of the barrel. However, an alternative, fully extended starting structure for the Nt region gave a height of 59 Å, very similar to those of the truncated systems. In this structure the Nt regions do not interact with the barrel.

## DISCUSSION

Our previous implicit simulations showed the CAV1 8S complex taking a conical shape in a variety of membrane and aqueous environments, in contrast to the starting flat shape of the 8S complex in the cryo-EM structure. This conical shape was also observed in the present all-atom simulations, almost to the same extent as the implicit solvent simulations. This conformational change also occurs in recent all-atom simulations of the 8S complex in a POPC/cholesterol membrane ^13^. The height of the 8S complex in the final unpalmitoylated system of that work is 56.3 Å, similar to ours. The only simulation that could maintain the flat shape of 8S was one in vacuum. One possible explanation then for the discrepancy between simulations and cryoEM is that DDM creates an overly nonpolar environment that strengthens salt bridges and other polar interactions and forces the complex into a flat shape. The low temperature of the experiment (liquid nitrogen, about 77 K) could also favor the more compact flat shape. To elucidate the role of detergent, all-atom simulations in DDM are currently in progress.

Another possible factor is the first 48 residues, which are absent in the simulations while present, yet invisible, in the experiment. In the context of heterologous caveolae in E. coli, this region has been found to be dispensable for membrane deformation, but its absence leads to smaller (i.e. more curved) and more monodisperse caveolae ^21^. Another study found that the Nt region is required for the higher oligomerization of caveolin into 70S complexes ^22^. It is noteworthy that one third of the pathogenic mutations in caveolin 3 occur in this region ^23^, as is the E38X mutation in CAV1 that causes lipodystrophy ^24^. The length of the Nt region is also relevant to the difference between the three caveolins, as it becomes shorter going from CAV1 to CAV2 and to CAV3. In addition, the β isoform of CAV1 misses the first 31 residues ^25^. Heterooligomers of CAV1 and CAV2 are reported to form deeper, i.e. more curved, caveolae than CAV1 homooligomers ^26^. It is thus conceivable that the Nt region may play a regulatory role by adjusting the shape of the complex. All-atom simulations in a bilayer could be conducted to validate this result, although the lack of a reliable structure is an impediment.

Regarding the way the CAV1 8S complex is embedded into a lipid bilayer, our results support the deep embedding found by other groups ^13,14^. The underside of the complex is almost completely hydrophobic. When it is placed on top of a bilayer, the zwitterionic phospholipid headgroups strongly attract water to hydrate themselves and lipids get inverted to solvate the caveolin complex. This results in reverse micelle formation and a bulge on the distal leaflet. More reasonable structures were obtained when the 8S complex started in the middle of the bilayer. The transformation to conical shape resulted in pulling the distal leaflet away from its original, flat position. This type of membrane deformation is closer to our expectation of positive membrane curvature generation. However, the lipid monolayer covering the 8S underside is not uniform but exhibits defects. The shape change of the protein and the distal leaflet call for some adjustments in the recent analytical model of curvature generation (Barnoy, Parton, and Kozlov 2024).

Liebl and Voth recently reported μs-long all-atom simulations of the 8S complex in 70:30 POPC/cholesterol membrane ^13^. They observed some extraction of cholesterol from the bilayer and then used metadynamics to specifically enhance cholesterol extraction. Here, in the absence of cholesterol we also see extraction of POPC. More work is needed to characterize the preference, if any, of cholesterol. The membrane structure in this work exhibits smaller defects than ours, which could be due to the presence of cholesterol. This might be a possible mechanism for cholesterol enrichment in caveolae, i.e. the small size of cholesterol may be able to better accommodate the deformations that the 8S imposes on the membrane. More work is needed to clarify this issue, including simulations with different lipid compositions. Coarse-grained simulations found that the 8S complex (largely constrained at the cryo-EM conformation) bends substantially the membrane ^14^, but this bending may be exaggerated ^13^.

Future studies could start from better initial system builds, for example with the protein already conical and the beta barrel filled with lipid. In addition, the periodic boundary conditions normally used impede the characterization of membrane curvature generation. This could be rectified by using noncontinuous bilayer patches, like bicelles and nanodisks. The effect of palmitoylation at residues 133,143, and 156 has already started to be explored ^13^and more could be done in the future. While palmitoylation is not necessary for the localization of CAV1 to caveolae, it is known to stabilize oligomers ^27,28^.

## Acknowledgements

This work was supported by the National Science Foundation (MCB 1855942). We thank Liebl and Voth for sharing the coordinates of their POPC/cholesterol bilayer simulations. Anton computer time was provided by the Pittsburgh Supercomputing Center through grant R01GM116961 from the NIH. The Anton machine at PSC was generously made available by D.E. Shaw Research, and computer time was provided by the National Center for Multiscale Modeling of Biological Systems through grant number P41GM103712-S1 from the NIH and Pittsburgh Supercomputing Center.

## TOC graphic

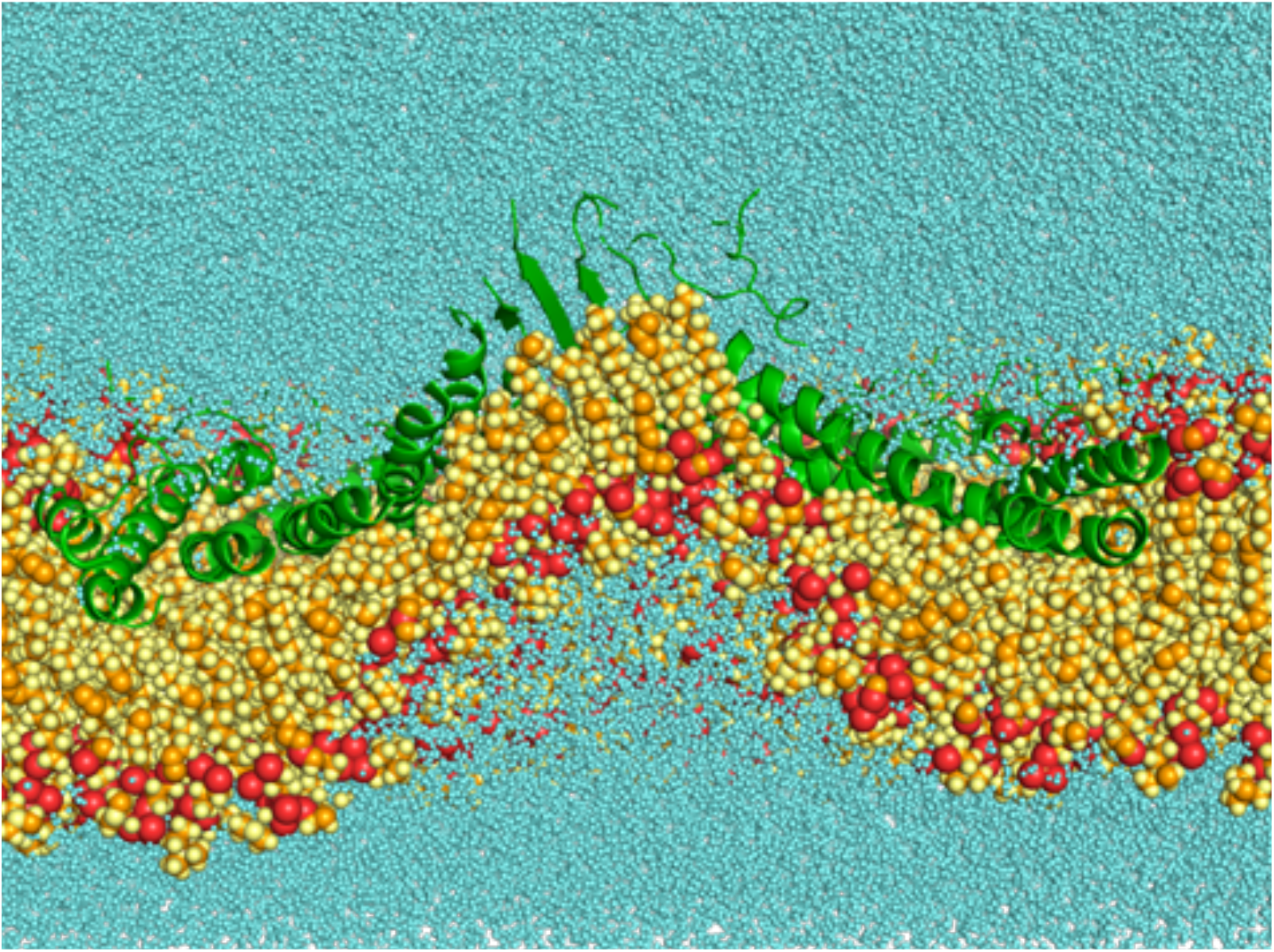

